# Thalamocortical networks involved in Pusher Syndrome

**DOI:** 10.1101/2022.10.12.511887

**Authors:** Hannah Rosenzopf, Julian Klingbeil, Max Wawrzyniak, Lisa Röhrig, Christoph Sperber, Dorothee Saur, Hans-Otto Karnath

## Abstract

Indirect quantification of functional and structural disconnection increases the knowledge derived from focal brain lesions by inferring subsequent brain network damage from the respective lesion. We applied both measures to a sample of 124 stroke patients to investigate brain disconnection in pusher syndrome – a disorder characterized by a disturbed perception of one’s own upright body posture. Our results suggest a hub-like function of the posterior and lateral portions of the thalamus in the perception of one’s own postural upright and identified dysfunction in a thalamo-cortical network as one likely cause of pusher syndrome. Lesion network-symptom-mapping investigating functional disconnection indicated cortical diaschisis in cerebellar, frontal, parietal, occipital, and temporal areas in patients with thalamic lesions suffering from pusher syndrome, but there was no evidence for functional diaschisis in cortical stroke and no evidence for the convergence of thalamic and cortical lesions onto a common functional network. Structural disconnectivity mapping identified several thalamo-cortical disconnections. Many of the thalamic and cortical regions disconnected by lesions that lead to pusher behavior correspond to previously reported lesion sites associated with pusher syndrome. Thus, while the presence of both, isolated thalamic and cortical lesions in context with pusher behavior has been reported previously and led to the conclusion that the correct estimation of one’s own postural upright might depend on a thalamo-cortical network, our analyses offer the first evidence for a direct thalamo-cortical (or cortico-thalamic) interconnection and, more importantly, shed light on the location of the respective thalamo-cortical disconnections.

## 1 Introduction

Stroke patients with ‘pusher syndrome’ (also termed ‘[contraversive] lateropulsion’ or ‘contraversive pushing’), a disorder occasionally accompanying hemiparesis, experience their body as oriented upright when it is markedly tilted to the ipsilesional side, indicating disturbed perception of upright body posture.^1^ They use the unaffected limb to actively push their body towards the paretic side, thus impairing their own balance and increasing the risk of falling in the direction of paresis.^2,3^

The neuronal correlates of the syndrome appear to be located in both thalamic^4–8^ and cortical regions.^6–11^ of both hemispheres. Thalamic locations include the ventral posterior and lateral posterior nuclei of the posterolateral thalamus.^4,5^ Cortical associations were reported for the insula,^6,8,10^ operculum,^6,8,10^ inferior frontal gyrus,^6^ superior temporal gyrus,^8^ precentral gyrus,^6^ dorsal antero-medial cortex,^11^ and postcentral gyrus,^6,9,10^ the latter extending towards the supramarginal gyrus.^9^ Alterations were further found in white matter structures, including the internal capsule,^4,6,8^ occipitofrontal fasciculus,^6^ uncinate fasciculus,^6^ and in the pathway between the vestibular nucleus and the parieto-insular vestibular cortex.^12^ Both, isolated cortical lesions^6–11^ and isolated thalamic lesions^4–8^ were found to cause pusher syndrome, which could indicate that the perception of one’s own postural upright relies on a brain network with at least two nodes. In addition to the local damage, stroke is known to result in functional deficits remote from the lesion. Von Monakow,^13^ who first described this phenomenon termed “diaschisis”, reasoned that it might be caused by collateral damage to connections between the lesion site and the respective dysfunctional areas. His hypothesis has since found support from modern neuroimaging methods.^14^ Pairwise functional connectivity has been found to generally reflect structural connectivity,^15,16^ although strong pairwise functional connections have also been reported between distant regions that do not appear to be directly linked by white matter.^15,17^

With the introduction of lesion network mapping, Boes and colleagues^18^ suggested an indirect method to investigate diaschisis in patients with focal brain lesions. They used normative connectome data from resting-state functional MRI of healthy participants to identify functional networks potentially affected by diaschisis. The lesion site is used as a seed for the functional connectivity analysis under the assumption that regions with high normative connectivity to the lesion are vulnerable to diaschisis.^19^ When the resulting lesion network maps are statistically compared between patients discordant for the symptom of interest, the method has been termed lesion network-symptom-mapping (LNSM).^20^ An alternative approach to investigate diaschisis is the indirect quantification of structural disconnection. Here, the structural lesion connectome is created by combining a patient’s topographical lesion data with normative white matter data to determine which grey matter parcels are most likely disconnected as a consequence of the respective lesion.^21,22^ Both functional and structural disconnectivity analyses have served to understand or predict several neuropsychological deficits and movement disorders secondary to stroke.^18,20,23^

Both methods have their individual advantages and shortcomings. Indirect measures of functional disconnection provide information on large-scale brain networks and can also account for diaschisis across multisynaptic pathways unlike directly and indirectly quantified structural disconnection. However, indirect functional disconnection measures appear to have a smaller predictive power on lesion-induced network dysfunctions than their structural counterparts.^29,30^ Despite their greater predictive power, structural disconnection tractograms are based on do not allow inference of the directionality of an uncovered disconnection.^31^ Further, when assessing the structural disconnectome, it is recommended to only consider streamlines which end in both parcels, to avoid a hardly interpretable overabundant amount of data.^22^ As a consequence, this method only documents the first grey matter parcel that is disconnected by the lesion and does not account for indirect damages further along in the network. Although both methods cannot be compared one-to-one, they offer insights into different aspects of the disrupted connectivity, and thus results may complement each other.

We attempted to, first, achieve a holistic view of the altered connectivity in pusher syndrome by investigating both types of connectivity and, second, to profit from the advantages of both approaches and compensate for their respective shortcomings, by applying both functional^18^ and structural disconnection measures.^22^ We utilized these methodologies in the largest cohort of patients reported so far with pusher syndrome characterized by a validated behavioural scale for pushing behavior (Scale of Contraversive Pushing [SCP]^1,2^). We hypothesized to identify a brain network affected by thalamic and cortical stroke through direct lesion and indirect diaschisis effects.

## 2 Material and methods

### 2.1 Patients and behavioral testing

Lesion masks and behavioral data were taken from three previous studies on the neural correlates of pusher syndrome by our group at the Neurology Department of the University Hospital in Tuebingen,^4,5,10^ detailed demographic tables can be drawn from the respective papers. In total, 131 patients with a single stroke event were examined. The first study^4^ included 23 patients consecutively admitted over two years with a circumscribed unilateral lesion and contraversive pushing (*Pusher+*) and 23 stroke patients admitted in the same period who did not exhibit contraversive pushing (*Pusher-*) but showed comparable age, lesion etiology and ratio of hemiparesis, spatial neglect and aphasia. The second study^5^ recruited 40 consecutive patients with thalamic strokes over two years and identified 14 *Pusher+* patients. The third study^10^ prospectively investigated 21 *Pusher+* that had suffered an acute stroke with unilateral cortical lesions sparing the thalamus. They had been consecutively admitted over 6 years. Two groups of control patients without pusher syndrome – 12 subjects with acute left-sided and 12 with acute right-sided cortical lesions sparing the thalamus – were matched to the groups of pusher patients. Seven patients (three *Pusher+* patients) from the latest study had already been included in one of the previous ones and were only included once in our final sample (n = 124). The studies had been approved by the Ethics Committee of the Medical Faculty at the University of Tübingen and all patients gave their written informed consent.

Pusher symptoms were examined with the standardized ‘Scale for Contraversive Pushing’ (SCP),^1,2^ which is based on Davies’ criteria.^32^ The SCP assesses a) the symmetry of patients’ spontaneous posture, b) arm and/or leg use as a means to increase contact with the ground and c) resistance to passive posture correction (all three subscales were measured for both sitting and standing). Contraversive pushing was diagnosed when for each subscale a score of at least 1 (out of 2, sitting plus standing) was measured.

For the LNSM, apart from analyzing the whole cohort, we also applied analyses after dividing patients with isolated cortical lesions and isolated thalamic lesions. These criteria were evaluated based on an overlay of the individual lesion masks with the AICHA brain atlas.^33^ This also resulted in the exclusion of nine patients who had both thalamic and cortical lesions; the subgroups then consisted of 72 patients with cortical stroke (31 *Pusher+* patients, Fig. 1C and D) and 43 patients with thalamic stroke (17 *Pusher+* patients, Fig, 1E and F). The subdivision into isolated thalamic and isolated cortical strokes was based on the assumption that diaschisis effects might differ between these groups. Indirect quantification of structural disconnectivity, on the other hand, is a procedure comparable to VLSM, with the difference that, instead of finding common grey matter substrates, the analysis yields uncovering commonly disconnected fiber tracts, based on patients’ lesions. When investigating unilateral lesions in opposing hemispheres in a single analysis, symmetric lesions of both hemispheres would act as opponents in the statistical analysis and corresponding effects might become distorted or remain undiscovered, i.e. one would run the risk of diluting existing effects. It has therefore become best practice to separate patients according to the lesioned hemisphere.^8,34,35^ In contrast to VLSM, indirect quantification of structural disconnectivity can, to some degree, also account for interhemispheric disconnections (a disconnection between one left and one right hemispheric region will be identified, no matter in which hemisphere the lesion occurs); since no gold standard has to date been established, we conducted both an analysis regardless of hemisphere and two analyses on patients separated by lesion side. The latter resulted in two samples of 49 patients with left hemispheric (21 *Pusher+* patients) and 75 patients with right hemispheric (34 *Pusher+* patients) lesions. In LNSM, on the other hand, shortcomings based on the pooling of right- and left hemispheric lesions are not expected to be pronounced, i.a. because lesion connectivity maps based on resting-state functional connectivity are highly symmetric.^36^ Accordingly, we are not aware of any LNSM study that separated left and right hemisphere lesions.

**Figure 1.**
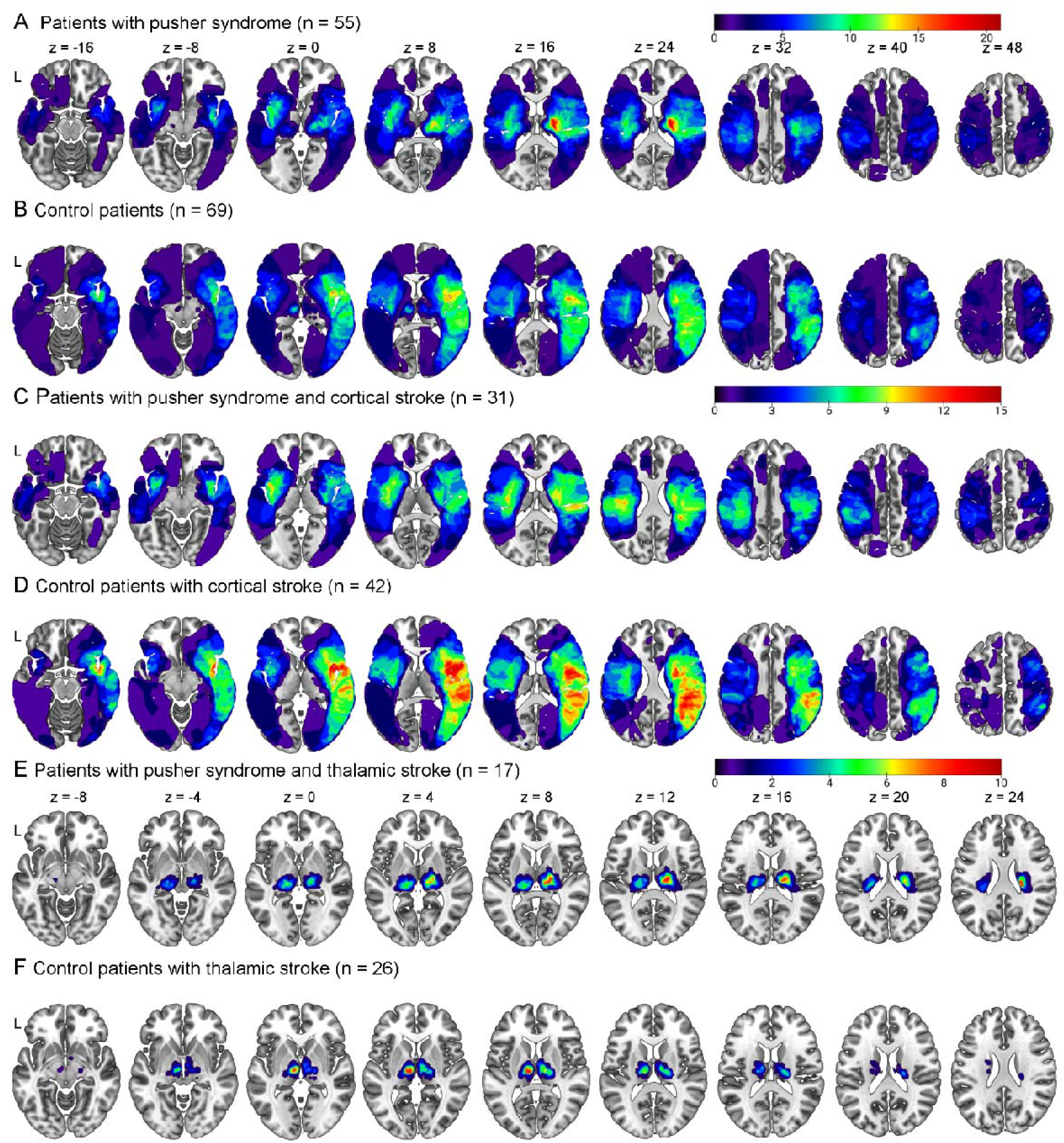
Lesion overlay. (A) Lesion overlay of all 55 stroke patients with pusher syndrome, (B) all 69 patients from the control cohort, (C) 31 patients with cortical lesions sparing the thalamus and pusher syndrome, (D) 41 patients with cortical lesions sparing the thalamus without pusher syndrome, (E) 17 patients with thalamic lesions sparing the cortex and pusher syndrome and (F) 26 patients with thalamic lesions sparing the cortex without pusher syndrome. Color indicates number of lesions. Axial coordinates refer to MNI space; L = left. Illustrations were rendered with MRIcroGL (https://www.nitrc.org/projects/mricrogl).

### 2.2 Lesion masks

Original lesion masks from the three previous studies^4,5,10^ were mapped onto template MRI scans in MNI152 (Montreal Neurological Institute) space. To 3-dimensionally reconstruct the lesions, we used these original lesion maps, expanded them along the z-axis by 4 or 5 mm and folded the resulting images with a Gaussian smoothing kernel with full width at half maximum of 4/4/10 mm (x/y/z). The resulting image was binarized at a threshold of ≥ 0.45. The resulting binary lesion maps (Fig. 1) were then used for LNSM, indirectly quantified structural disconnection mapping, and voxel-based lesion-symptom-mapping (VLSM, see supplementary material) analyses.

### 2.3 Magnetic resonance imaging and preprocessing

For LNSM, resting-state fMRI data from healthy young participants (n = 100) made available by the human connectome project^37^ were used to calculate lesion network maps with Statistical Parametric Mapping (SPM12, v7487, Wellcome Trust Centre for Neuroimaging, London) and Matlab tools programmed specifically for this purpose (R2019a, MathWorks). Functional data sets consisted of two resting-state sessions (left-right and right-left phase encoding, gradient-echo EPI sequence, TR 720 ms, resolution 2 mm isotropic, 15 minutes). All downloaded images had already been ‘minimally preprocessed’^38^ and were further convolved with an isotropic Gaussian smoothing kernel (full width at half maximum: 5 mm). A multiple regression approach was applied to remove signal variance over time explained by nuisance variables (motion parameters, mean white matter, CSF, global signal). A band-pass filter was applied on the residual BOLD time series (0.01 - 0.08 Hz). Two datasets from controls were excluded due to heavy in-scanner motion with less than 10 minutes of restingstate fMRI remaining in one session after motion scrubbing (> 0.5 mm of frame-wise displacement lead to scrubbing). The individual brain lesions (restricted to regions with a grey matter tissue probability > 10%) were defined as regions of interest. The first eigenvariate of the time series of all voxels within that ROI was the representative BOLD time series. We then calculated lesion network maps based on functional connectivity (FC) (i.e., Fisher-transformed Pearson correlation coefficients) between ROI time series and the time series of all other brain voxels. Separate connectivity maps were generated for the right-left and left-right phase encoding session, but to receive one lesion network for each patient, both connectivity maps from all 98 remaining controls were averaged.

#### Lesion network-symptom-mapping

LNSM was carried out as described before^20^ with non-parametric permutation testing.^40^ To reveal differences in lesion networks between patient groups (e.g., *Pusher+* vs. *Pusher-*), two-sample t-tests were conducted in every voxel. The null distribution of resulting t-scores was obtained by repetition of the statistical test with randomly assigned group labels 5000 times. The initial test result (with correct group assignments) was then thresholded at a t-value corresponding to p(FWE) < 0.05 at cluster level. Anatomical labeling was performed with the AICHA atlas^33^ or the AAL2 atlas^41^ for the cerebellum.

#### Indirect quantification of structural disconnection

Individual damage to structural brain networks was indirectly assessed by creating a parcelwise disconnection matrix with the lesion quantification toolkit (LQT)^22^; operated on MATLAB R2020a (Mathworks). To get an impression to what extent the use of different parcellations affects the resulting disconnections, we used two different thalamus atlas parcellations, the Oxford thalamic connectivity atlas^42,43^ and a more finely grained connectivity-based thalamus parcellation by Kumar and colleagues.^44^ We combined each with the cortical parcellation provided by Desikan and colleagues.^45^ In a separate run for each of the two brain parcellations, we used the LQT to obtain a disconnectivity matrix for both parcellations for each patient. To do so, we first used the LQT to combine the respective brain parcellation with the HCP-842 streamline tractography atlas^46^ to create a structural connectivity matrix representing the healthy human connectome. Each lesion was then individually integrated as a region of interest in the tractography atlas. A filter identified streamlines, which (i) were disconnected by the lesion (i.e., crossed the binary lesion map) and (ii) terminate in an atlas region on both ends (i.e., represent a connection between two atlas regions) defined by the atlas parcellation. The resulting matrix containing absolute disconnection counts between each pair of regions is then related to the original connectivity matrix to create a final matrix, where each cell represents the percentage of disconnected fiber bundles between two atlas regions.

We tested the resulting disconnectivity matrices for disconnection-deficit associations with a mass-univariate analysis using logistic regression with custom scripts in Matlab. Within the symmetric disconnectivity matrices, we removed the diagonal and all redundant elements below it. Next, we removed region-to-region connections that were damaged in less than 10 patients. This procedure mirrored standard procedures in VLSM^47^ and prevented connections with an uninformative low pathological variance to affect statistical results. The remaining 549 region-to-region connections in the parcellation with the Oxford thalamic connectivity atlas^42,43^ and 626 in the parcellation with the thalamus atlas by Kumar and colleagues^44^. were subjected to the statistical analysis. For each of the remaining region-to-region connections, a logistic regression was run to model the binary pusher diagnosis from continuous disconnection scores. Statistical significance was assessed by maximum statistic permutation^40^ with 10000 random permutations and a one-sided statistical significance level of p < 0.05. This serves as an approximately exact family-wise error correction for multiple comparisons. The same procedure was also conducted with data separated by lesion side (right versus left hemispheric lesions).

## 3 Results

### Lesion network-symptom-mapping

With LNSM we found no significant differences between *Pusher+* and *Pusher-* patients for the whole cohort and the subgroup with isolated cortical lesions (Fig. 2A and 2B, respectively). In the subgroup of patients with isolated thalamic lesions, however, we identified a significant difference in normative lesion connectivity to seven different clusters (Fig. 2C). Five clusters showed significantly higher normative lesion connectivity in *Pusher+* patients (‘right inferior parietal’, ‘right anterior temporomesial’, ‘right posterior temporomesial’, ‘left and right lateral occipital cortex and precuneus’) and two clusters showed significantly lower lesion connectivity (‘bilateral cerebellum and vermis’, ‘left paracingulate and cingulate gyri’).

**Figure 2.**
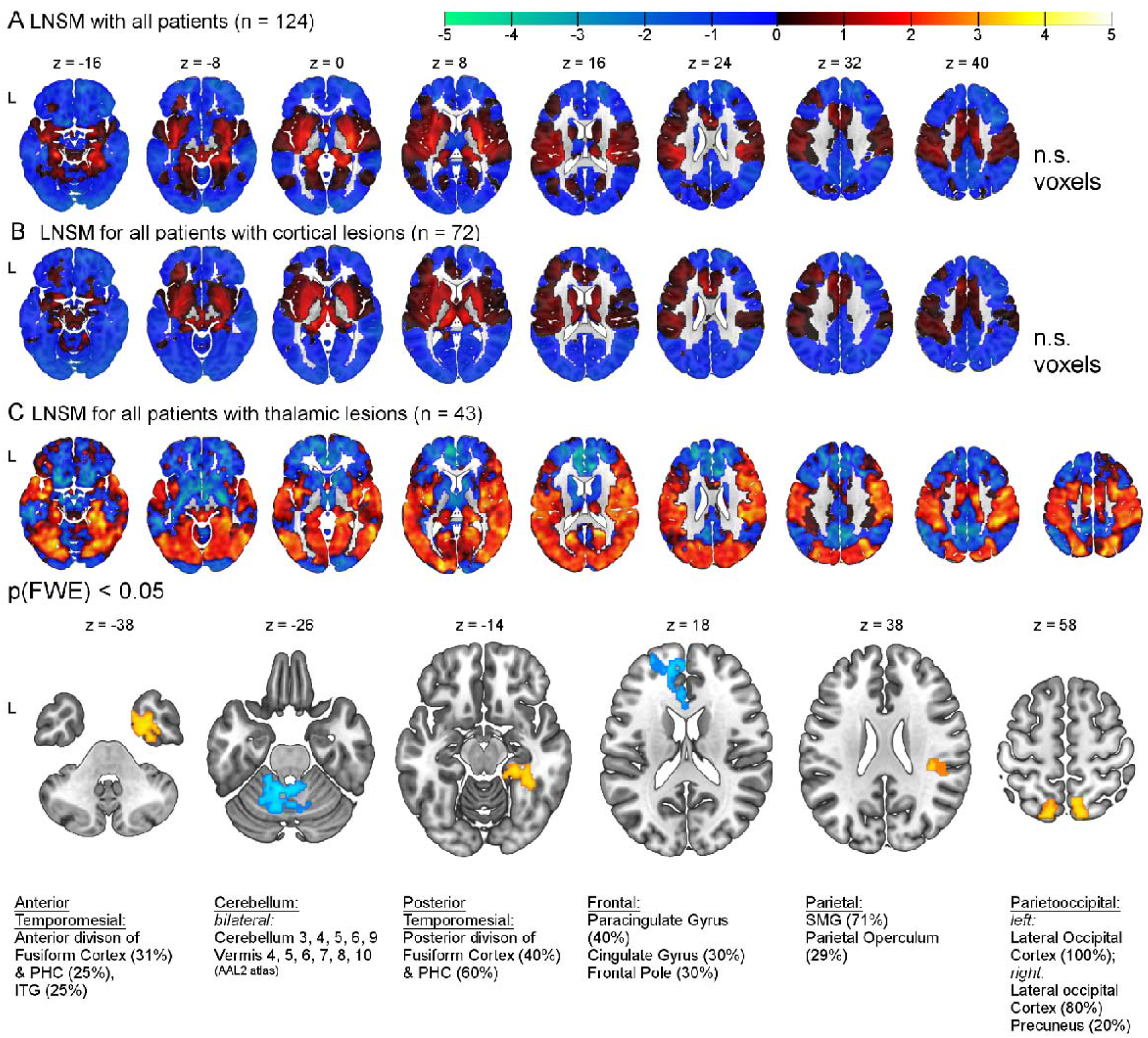
Lesion networks associated with pusher syndrome. Statistical inference was based on a random permutation test thresholded at p(FWE) < 0.05 at the cluster-level. The LNSM found no significant results for the whole cohort (A) and the subgroup of patients with cortical lesions that spare the thalamus (B). The LNSM revealed significant differences in the subgroup of patients with thalamic lesions that spare the cortex (C): In this cohort, higher or lower lesion connectivity to several cortical regions was associated with the pusher syndrome. Abbreviations: FWE = family-wise error; ITG = inferior temporal gyrus; L = left; PHC = parahippocampal cortex; SMG = supramarginal gyrus. Scale: t-values; coordinates in MNI space.

In a further analysis, we restricted the LNSM in patients with isolated cortical stroke to those thalamic regions where lesions cause pusher syndrome. This was achieved with the use of a binary mask that included all thalamic voxels where at least one *Pusher+* patient in the whole cohort has a lesion (see Fig. 3A, in beige). We also analyzed patients with isolated thalamic lesions using a mask for cortical lesion in *Pusher+* (see Fig. 3A, in beige). LNSM was applied as before, but the use of these masks results in a small volume correction limiting the number of statistical comparisons. Still, no significant difference between *Pusher+* and *Pusher*patients was identified in the thalamic mask for the subgroup of patients with cortical stroke (Fig. 3A). In contrast, for the subgroup of patients with thalamic lesions, five clusters were identified (Fig. 3B). Four clusters showed significantly higher normative lesion connectivity in *Pusher+* patients (‘left and right inferior parietal’, ‘right middle temporal’, ‘left lateral occipital cortex’) and one cluster showed significantly lower lesion connectivity (‘left paracingulate and cingulate gyri’).

**Figure 3.**
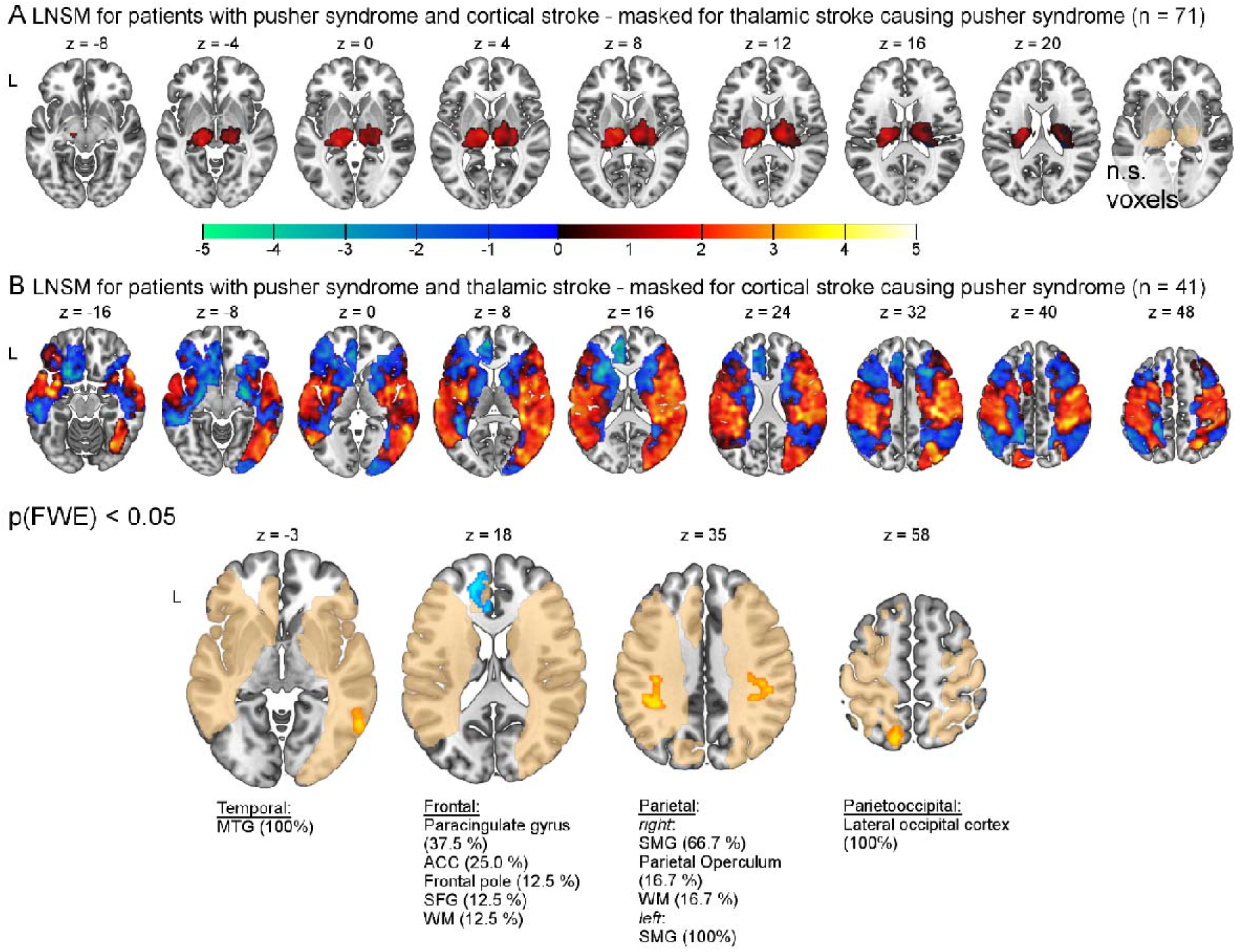
Lesion networks associated with pusher syndrome. Applied brain masks illustrated in beige show all (A) thalamic or (B) cortical voxels, respectively, damaged in at least one pusher patient. Statistical inference was based on a random permutation test thresholded at p(FWE) < 0.05 at the cluster-level. (A) With LNSM no evidence for diaschisis in the thalamus were found when the analysis was restricted the effect of cortical lesions on thalamic lesions known to cause pusher syndrome when damaged. (B) However, LNSM revealed significant differences in the subgroup of patients with thalamic lesions when the analyses were restricted to cortical regions also known to cause pusher syndrome. Higher or lower lesion connectivity to a total of five regions was associated with the presence of pusher syndrome. Abbreviations: ACC = anterior cingulate cortex; FWE = family-wise error; L = left; MTG = middle temporal gyrus = MTG; PHC = parahippocampal cortex; SFG = superior frontal gyrus; SMG = supramarginal gyrus; WM = white matter. Scale: t-values; coordinates in MNI space.

### Indirect quantification of structural disconnection

Indirectly quantified structural disconnection-symptom mapping based on the indirectly assessed disconnection matrices on the complete sample revealed no significant connections for the parcellation with the Oxford thalamic connectivity atlas^42,43^ after correction for multiple comparisons. With the parcellation based on the thalamus atlas by Kumar and colleagues^44^, one single connection was found to be significant after family-wise error correction. Disconnection between parcel number 13 and the right precentral gyrus was significantly associated with the pusher syndrome at an uncorrected p = 0.0015 (corrected p < 0.05). Region 13 in the right hemisphere shows the greatest overlap with the medial geniculate nucleus of the posterior group, followed by the ventroposterior complex in the lateral group and, to a lesser extent, several other posterior nuclei^44^. The equivalent connection in the left hemisphere was not found to be significant (uncorrected p = 0.57).

After separating lesion masks based on the affected hemisphere, pusher syndrome in association with left hemispheric lesions, the left Kumar thalamus region 15 (‘posterior group’)^44^ displayed significant disconnections to the left banks of the superior temporal sulcus and the left superior temporal gyrus. Analysis with the Oxford thalamic connectivity atlas, on the other hand, did not yield any significant left hemispheric disconnections. Concerning right hemispheric lesions and the Kumar parcellation, four disconnections were found between thalamus region 1 (‘posterior group’)^44^ and the postcentral gyrus, thalamus region 8 (‘lateral group’)^44^ and the paracentral lobule, and thalamus region 13 (‘posterior group’)^44^ and both the pre- and postcentral gyrus. Based on the Oxford thalamic parcellation, we found four significant right hemispheric disconnections: the postcentral gyrus was found to be disconnected from two thalamus parcels primarily reported for their primary and premotor projections^42,43^; the latter thalamic parcel, which is supposed to be responsible for premotor projections,^42,43^ showed two further disconnections to the banks of the temporal sulcus and the inferior temporal gyrus. All disconnections are illustrated in figure 4.

**Figure 4.**
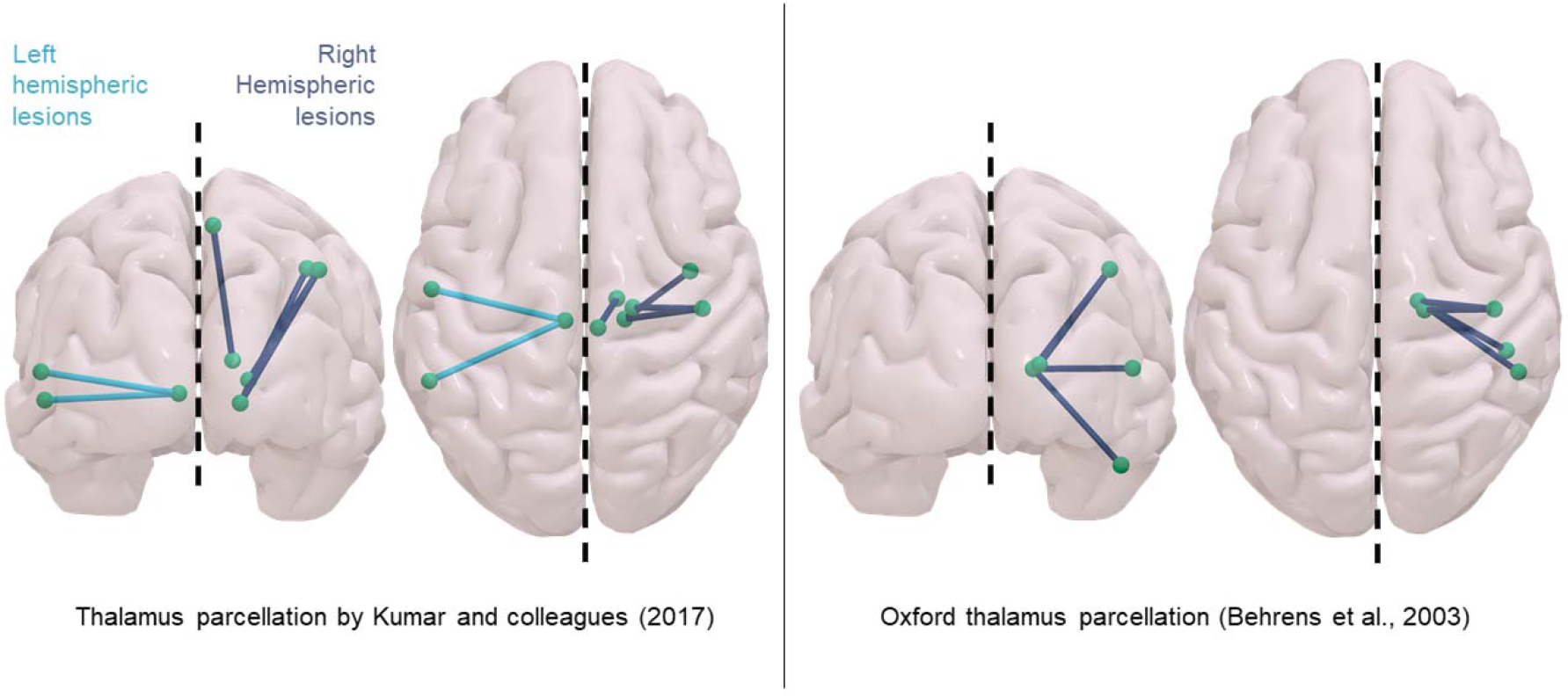
Structural ROI to ROI disconnections associated with pusher syndrome. Significant (p < 0.05) disconnections obtained from the indirect quantification of structural disconnectivity with the LQT^22^ in the sub-samples separated by lesioned hemisphere. Illustrations were generated with Surf Ice (https://www.nitrc.org/projects/surfice/). Nodes and edges for 3D reconstruction of the respective disconnections can be found in the online materials.

## 4 Discussion

In the present study, differences between stroke patients with and without pusher syndrome were investigated with indirect measures of functional and structural brain disconnectivity. The analyses were based on lesions and behavioral data from 124 stroke patients from three earlier localisatory studies,^4,5,10^ which had demonstrated that both cortical and thalamic lesions in the left and right hemisphere may cause pusher syndrome, indicating a brain network with at least two nodes. The LNSM performed in the present study identified cortical diaschisis after thalamic lesions but neither evidence for thalamic diaschisis after cortical lesions nor shared functional disconnection in both thalamic and cortical stroke. Our analyses of indirectly quantified structural disconnection uncovered exclusively thalamo-cortical disconnectivity in association with pusher syndrome. In the following, these functional and structural aspects of our investigation are discussed in detail.

### 4.1. Functional disconnection

With LNSM, we found no evidence for the convergence of thalamic and cortical lesions onto a common functional network in pusher syndrome. We used two further analyses to determine if pusher syndrome caused by thalamic lesions might result from cortical diaschisis or if pusher syndrome caused by cortical lesions might be a consequence of thalamic diaschisis. We found no evidence for functional diaschisis in the thalamus caused by cortical stroke (Fig. 2B). This was even true when the analyses were restricted to thalamic regions known to cause pusher syndrome – which simultaneously amounts to a small volume correction and thus further underscores the absence of evidence for diaschisis effects in the thalamus as a result of cortical stroke in pusher syndrome (Fig. 3A).

However, we identified diaschisis effects in several cortical regions associated with lesions in the thalamus that cause pusher syndrome. These included regions with higher lesion connectivity (right inferior parietal areas, right middle temporal areas, the bilateral lateral occipital cortex and precuneus) and regions with lower lesion connectivity (left paracingulate and cingulate gyri, bilateral cerebellar regions) (Fig. 2C). When restricting the analysis to cortical regions known to cause pusher syndrome in our cohort, we observed the inferior parietal regions bilaterally (Fig. 3B). Based on the identification of thalamic damage in pusher syndrome and the known posterior thalamic connections with the occipital, parietal, and temporal cortices,^42,43^ dysfunction or metabolic abnormalities in the temporal, parietal and occipital cortex had been expected to contribute to the pusher syndrome.^5^

The inferior parietal region, where diaschisis was associated with the pusher syndrome in patients with thalamic lesions, has already been found to be associated with pusher syndrome in three independent lesion studies^7,9,48^ and one previous study based on a part of the current data set.^10^ It has been described to be the most relevant lesion location in pusher syndrome^9^ and was hypothesized to be a sensory integration area where an egocentric reference system for vertical upright posture is created.^48,49^

The temporal areas, where diaschisis was associated with the pusher syndrome in patients with thalamic lesions, are part of the multimodal vestibular cortex that spans a large temporoparietal perisylvian area.^50^ Apart from vestibular cortex, we also found an association of the pusher syndrome and diaschisis in the lateral occipital cortex, which might interfere with the integration of visual information. Moreover, the identification of the anterior and posterior temporomesial regions might also be related to multisensory integration. It has been shown that stance and visually guided locomotion involve the posterior hippocampal formation while vestibular signals enter the anterior part of the hippocampal formation.^51^

### 4.2 Structural disconnection

All structural disconnections found with pusher syndrome in the present study consisted of streamlines between one thalamic and one cortical grey matter parcel. Many of these thalamic and cortical parcels correspond to previously reported lesion locations associated with the disorder. Thalamic projections seem to be particularly affected in its posterior and lateral portions, which is in line with previous work by Karnath and colleagues.^4,5^ It is of note that the patients analyzed in the aforementioned studies were also part of the present sample, which of course presupposes overlapping results to some degree. However, the fact that these areas persisted in a larger sample reinforces their association with pusher syndrome.

On the cortical level, the uncovered structural disconnection involving the precentral gyrus, postcentral gyrus, and superior temporal gyrus corresponded with previous reports of grey matter damage associated with pusher syndrome,^6,10^ also in studies with patient samples independent from those studied here.^8,9^ The prominent role of the latter two regions in the perception of the postural upright is further stressed by a study that applied cortical electrical stimulation in epileptic patients.^52^ Kahane and colleagues’ stimulation of the postcentral and the superior temporal gyri elicited the perception of displacement/tilt in the roll plane. The suggested relevance of the postcentral and the temporal gyri are further in line with a second pathway proposed by Mittelstaedt^53^ in the context of human graviception, namely the processing of graviceptional information transported from the abdomen via the afferent sympathetic and parasympathetic nervous systems. In this process, Mittelstaedt^53^ suggested a potential afferent involvement of the vagal nerve. In accordance, activity not only in bilateral portions of the postcentral gyrus but also in posterior portions of the right temporal gyrus poses a reported consequence of such activation.^54^

Interestingly, the paracentral lobule and the inferior temporal gyrus are, to our knowledge, observed for the first time in context with pusher behavior. In the approach applied in the present study, lesions affecting either end of a damaged connection contribute collectively to its discovery, whereas VLSM □ used in previous studies □ detects only directly lesioned areas. In a patient cohort where both ends are damaged at a moderate rate, VLSM might therefore fail to detect either of them. Under similar circumstances, methods quantifying structural disconnection have better chances to detect both grey matter nodes (by identifying the related disconnected fiber tract), thanks to their pooled statistical contribution.^55^ Measures of structural disconnectivity have previously been found to have a predictive power comparable to focal lesions for several behavioral deficits^29^ and fare even better than grey matter damage in explaining brain network dysfunction after stroke.^30^ These perks might have contributed to the uncovering of so far unknown areas associated with pusher syndrome.

### 4.3 The importance of thalamo-cortical disconnection

Analyses of both left and right hemispheric lesions demonstrated exclusively thalamo-cortical structural disconnections in association with pusher syndrome. While the presence of isolated thalamic and isolated cortical lesions in the context of pusher behavior has been reported previously and led to the conclusion that correct estimation of one’s own postural upright might depend on a thalamo-cortical network, our analyses offer the first evidence for a direct thalamo-cortical (or cortico-thalamic) interconnection and shed light on the location of the respective thalamo-cortical disconnections.

The present results suggest a hub-like function of the posterior and lateral portions of the thalamus in the correct assessment of one’s own postural upright. The lack of significant cortico-cortical (and possibly also thalamo-thalamic) disconnections does not necessarily imply that their role is negligible. Their underrepresentation in comparison to thalamo-cortical disconnections might have masked effects on connections located exclusively in the thalamus (only possible in thalamic lesions) and exclusively in the cortex (possible only in cortical lesions), in a similar manner as right hemispheric lesions in VLSM weaken left-hemispheric ones and vice-versa. Comparing the results obtained from the complete sample to those of the subsamples separated by lesioned hemisphere indicates that a lack of appropriate subsampling as in the first case might have equally led to the underpowering of unilateral thalamo-cortical disconnections.

While the importance of disconnection between the thalamus and cortex was stressed by the indirect quantification of structural disconnectivity, the method fails to supply information about the directionality of these disconnections. However, LNSM found only thalamo-cortical but not cortico-thalamic diaschisis, which might imply thalamo-cortical rather than corticothalamic disconnection. The thalamus is a mediator of cortico-cortical communication which depends on higher-order thalamic relay nuclei with afferent and efferent connections to various cortical regions.^56^ Damage to a network hub such as the posterior thalamus is especially prone to cause widespread diaschisis. Our structural disconnection mapping provides evidence for the hub-like function of the posterior and lateral portions of the thalamus in the correct assessment of one’s own postural upright. Our functional disconnection mapping points towards distinctive mechanisms that cause pusher syndrome in cortical and thalamic stroke. The pusher syndrome in thalamic stroke patients might be a consequence of a failed multisensory integration because of cortical diaschisis after the loss of the central network hub. Meanwhile, pusher syndrome in cortical stroke might result from a failed multisensory integration brought about by the loss of critical network nodes sustaining one of the several necessary subfunctions. Nevertheless, future research is needed to support these hypotheses.

### 4.4 Limitations

There are several limitations to our analyses that need to be considered. First, apart from the disconnections involving the postcentral gyrus, structural disconnectivity results from the different parcellations varied considerably. Disconnection between the thalamus and the banks of the superior temporal sulcus was associated with pusher behavior in patients with lesions in both hemispheres. However, the left and right disconnections were identified by different atlas parcellations and seem to have slightly different thalamic origins. While Kumar’s asymmetrical parcellation might have masked results in the right hemisphere, the Oxford parcellation might have failed to uncover comparable results in both hemispheres because the left hemispheric sample was considerably smaller. Our results demonstrate the decisive role the choice of parcellation plays in successfully identifying neural correlates of pathologic behavior, while neither of the results can be classified as more or less accurate at this point. Further investigations are needed to evaluate which specific thalamo-cortical disconnections are more likely linked to pusher behavior.

Second, the fact that pusher syndrome is relatively rare and can be caused by left and right hemisphere stroke renders our sample rather unique. Most disorders originating from brain damage are (more) frequently observed and/or are predominantly lateralized, so previous structural disconnection studies focused on one hemisphere.^57–60^ But due to the more heterogeneous anatomy of pusher syndrome and the different prerequisites concerning the investigation of functional and structural disconnection, our sample had to be treated differently to obtain meaningful results (i.e., separation according to thalamic versus cortical lesion or separation according to left versus right hemisphere, respectively). While this is necessary for the interpretability of the individual analyses, it impedes a direct comparison of the results derived from the two different methods. Accordingly, our goal was *not* to directly compare both methods and their results. Much rather, we strived for a holistic view of alterations in the connectivity network by joining the complementary methods. Although the methods used cannot directly support or challenge each other’s’ results, due to the reported diversity between functional and structural connectivity,^15,17^ they both offer individual contributions to a better understanding of pusher syndrome as a network disorder.

Third, the inability to identify a common functional network associated with pusher syndrome in *both* thalamic and cortical lesions might be rooted in the limitations of LNSM. The ability of functional connectivity measures to identify the neural basis of higher cognitive functions has been demonstrated previously.^61^ Resting-state fMRI acquired in stroke patients could predict behavioral deficits in associative cognitive domains like verbal or visual memory better than lesion location. This was interpreted as evidence for an organization of these functions in large-scale networks. However, LNSM relies on an indirect measure of functional connectivity. While this approach was used to map the neural correlates of numerous neurological deficits,^18,23–25,62^ major limitations have been identified.^29,63,64^ The low dimensionality of lesion network maps might be the main reason why predictive modeling with lesion network data has failed so far.^29,64^ This problem is more pronounced when LNSM is based on large lesions that contain multiple discrete parcels of functionally heterogeneous tissue.^29,65^ This might also explain why, in our sample, only the analyses based on the comparatively small thalamic lesions identified diaschisis.

Finally, our study had to rely on the original lesion maps drawn onto predefined slices of an MNI-template which were then expanded along the z-axis to create lesion volumes, rather than using the clinical images themselves. This is in line with lesion network mapping studies from other groups,^18,23,24,66,67^ and it allowed us to compile the largest lesion dataset of pusher patients to date, albeit with reduced spatial accuracy of the lesion maps.

## Conclusions

The present results suggest a hub-like function of the posterior and lateral portions of the thalamus in the perception of one’s own postural upright. Based on the indirect quantification of both, functional and structural disconnection, we identify dysfunction in a thalamo-cortical network as one likely cause of pusher syndrome. Due to their effortless availability from stroke imaging, indirect measures of brain disconnection can contribute to a better understanding of the complex neural correlates of less common stroke symptoms. Still, future work in pusher syndrome should aim at validating our results with direct measures of functional and structural disconnection. Taken together, our findings contribute to an understanding of pusher syndrome as a thalamo-cortical network disorder.

## Funding

Julian Klingbeil and Dorothee Saur were supported by the German Research Foundation (SA 1723/5-1), Hans-Otto Karnath by the German Research Foundation (KA 1258/23-1).

## Notes

### Competing Interest Statement

The authors have declared no competing interest.

## References

1. Karnath HO, Ferber S, Dichgans J. The origin of contraversive pushing: Evidence for a second graviceptive system in humans. Neurology. 2000;55(9):1298–1304. doi:10.1212/WNL.55.9.1298

2. Karnath HO, Brötz D, Götz A. [Clinical symptoms, origin, and therapy of the “pusher syndrome”]. Nervenarzt. 2001;72(2):86–92. doi:10.1007/s001150050719

3. Karnath HO. Pusher syndrome - A frequent but little-known disturbance of body orientation perception. J Neurol. 2007;254(4):415–424. doi:10.1007/s00415-006-0341-6

4. Karnath HO, Ferber S, Dichgans J. The neural representation of postural control in humans. Proc Natl Acad Sci U S A. 2000;97(25):13931–13936. doi:10.1073/pnas.240279997

5. Karnath H-O, Johannsen L, Broetz D, Küker W. Posterior thalamic hemorrhage induces “pusher syndrome.” Neurology. 2005;65(10):1682. doi:10.1212/WNL.65.10.1682

6. Ticini LF, Klose U, Nägele T, Karnath HO. Perfusion imaging in pusher syndrome to investigate the neural substrates involved in controlling upright body position. PLoS One. 2009;4(5). doi:10.1371/journal.pone.0005737

7. Santos-Pontelli TEG, Pontes-Neto OM, Araujo DB de, Santos AC dos, Leite JP. Neuroimaging in stroke and non-stroke pusher patients. Arq Neuropsiquiatr. 2011;69(6):914–919. doi:10.1590/s0004-282x2011000700013

8. Baier B, Janzen J, Müller-Forell W, Fechir M, Müller N, Dieterich M. Pusher syndrome: Its cortical correlate. J Neurol. 2012;259(2):277–283. doi:10.1007/s00415-011-6173-z

9. Babyar SR, Smeragliuolo A, Albazron FM, Putrino D, Reding M, Boes AD. Lesion Localization of Poststroke Lateropulsion. Stroke. 2019;50(5):1067–1073. doi:10.1161/STROKEAHA.118.023445

10. Johannsen L, Broetz D, Naegele T, Karnath HO. “Pusher syndrome” following cortical lesions that spare the thalamus. J Neurol. 2006;253(4):455–463. doi:10.1007/s00415-005-0025-7

11. Karnath HO, Suchan J, Johannsen L. Pusher syndrome after ACA territory infarction. Eur J Neurol. 2008;15(8):84–85. doi:10.1111/j.1468-1331.2008.02187.x

12. Yeo SS, Jang SH, Oh S, Kwon JW. Role of diffusion tensor imaging in analyzing the neural connectivity of the parieto-insular vestibular cortex in pusher syndrome: As case report. Medicine (Baltimore). 2020;99(16):e19835. doi:10.1097/MD.0000000000019835

13. von Monakow, C. Die Lokalisation im Grosshirn und der Abbau der Funktion durch kortikale Herde. Bergmann; 1914.

14. Meyer JS, Obara K, Muramatsu K. Diaschisis. Neurol Res. 1993;15(6):362–366. doi:10.1080/01616412.1993.11740164

15. Greicius MD, Supekar K, Menon V, Dougherty RF. Resting-state functional connectivity reflects structural connectivity in the default mode network. Cereb Cortex. 2009; 19(1): 72–78. doi:10.1093/cercor/bhn059

16. Hermundstad AM, Bassett DS, Brown KS, et al. Structural foundations of resting-state and task-based functional connectivity in the human brain. Proc Natl Acad Sci US A. 2013;110(15):6169–6174. doi:10.1073/pnas.1219562110

17. Honey CJ, Sporns O, Cammoun L, et al. Predicting human resting-state functional connectivity from structural connectivity. Proc Natl Acad Sci U S A. 2009;106(6):2035–2040. doi:10.1073/pnas.0811168106

18. Boes AD, Prasad S, Liu H, et al. Network localization of neurological symptoms from focal brain lesions. Brain. 2015;138(10):3061–3075. doi:10.1093/brain/awv228

19. Fox MD. Mapping Symptoms to Brain Networks with the Human Connectome. N Engl J Med. 2018;379(23):2237–2245. doi:10.1056/nejmra1706158

20. Wawrzyniak M, Klingbeil J, Zeller D, Saur D, Classen J. The neuronal network involved in self-attribution of an artificial hand: A lesion network-symptom-mapping study. Neuroimage. 2018;166:317–324. doi:10.1016/j.neuroimage.2017.11.011

21. Foulon C, Cerliani L, Kinkingnéhun S, et al. Advanced lesion symptom mapping analyses and implementation as BCBtoolkit. Gigascience. 2018;7(3):1–17. doi:10.1093/gigascience/giy004

22. Griffis JC, Metcalf N V., Corbetta M, Shulman GL. Lesion Quantification Toolkit: A MATLAB software tool for estimating grey matter damage and white matter disconnections in patients with focal brain lesions. NeuroImage Clin. 2021;30:102639. doi:10.1016/j.nicl.2021.102639

23. Joutsa J, Horn A, Hsu J, Fox MD. Localizing parkinsonism based on focal brain lesions. Brain. 2018;141(8):2445–2456. doi:10.1093/brain/awy161

24. Darby RR, Laganiere S, Pascual-Leone A, Prasad S, Fox MD. Finding the imposter: Brain connectivity of lesions causing delusional misidentifications. Brain. 2017;140(2):497–507. doi:10.1093/brain/aww288

25. Padmanabhan JL, Cooke D, Joutsa J, et al. A human depression circuit derived from focal brain lesions. Biol Psychiatry. 2019;86(10):749–758. doi:doi:10.1016/j.biopsych.2019.07.023

26. Klingbeil J, Wawrzyniak M, Stockert A, et al. Pathological laughter and crying: Insights from lesion network-symptom-mapping. Brain. 2021;144(10):3264–3276. doi:10.1093/brain/awab224

27. Wiesen D, Karnath HO, Sperber C. Disconnection somewhere down the line: Multivariate lesion-symptom mapping of the line bisection error. Cortex. 2020;133:120–132. doi:10.1016/j.cortex.2020.09.012

28. Kuceyeski A, Navi BB, Kamel H, et al. Structural connectome disruption at baseline predicts 6-months post-stroke outcome. Hum Brain Mapp. 2016;37(7):2587–2601. doi:10.1002/hbm.23198

29. Salvalaggio A, de Filippo De Grazia M, Zorzi M, de Schotten MT, Corbetta M. Poststroke deficit prediction from lesion and indirect structural and functional disconnection. Brain. 2020;143(7):2173–2188. doi:10.1093/brain/awaa156

30. Griffis JC, Metcalf N V., Corbetta M, Shulman GL. Structural Disconnections Explain Brain Network Dysfunction after Stroke. Cell Rep. 2019;28(10):2527–2540.e9. doi:10.1016/j.celrep.2019.07.100

31. Mori S, Crain BJ, Chacko VP, Van Zijl PCM. Three-dimensional tracking of axonal projections in the brain by magnetic resonance imaging. Ann Neurol. 1999;45(2):265–269. doi: 10.1002/1531-8249(199902)45:2<265::AID-ANA21>3.0.CO;2-3

32. Davies P.M. Steps to follow: a guide to the treatment of adult hemiplegia. Springer; 1985.

33. Joliot M, Jobard G, Naveau M, et al. AICHA: An atlas of intrinsic connectivity of homotopic areas. J Neurosci Methods. 2015;254:46–59. doi:10.1016/j.jneumeth.2015.07.013

34. Frenkel-Toledo S, Fridberg G, Ofir S, et al. Lesion location impact on functional recovery of the hemiparetic upper limb. PLoS One. 2019;14(7):1–28. doi:10.1371/journal.pone.0219738

35. Frenkel-Toledo S, Ofir-Geva S, Mansano L, Granot O, Soroker N. Stroke Lesion Impact on Lower Limb Function. Front Hum Neurosci. 2021; 15(February): 1–11. doi:10.3389/fnhum.2021.592975

36. Raemaekers M, Schellekens W, Petridou N, Ramsey NF. Knowing left from right: asymmetric functional connectivity during resting state. Brain Struct Funct. 2018;223(4):1909–1922. doi:10.1007/s00429-017-1604-y

37. Yeo BTT, Krienen FM, Sepulcre J, et al. The organization of the human cerebral cortex estimated by intrinsic functional connectivity. J Neurophysiol. 2011;106(3):1125–1165. doi:10.1152/jn.00338.2011

38. Glasser MF, Sotiropoulos SN, Wilson JA, et al. The minimal preprocessing pipelines for the Human Connectome Project. Neuroimage. 2013;80:105–124. doi:10.1016/j.neuroimage.2013.04.127

39. Power JD, Barnes KA, Snyder AZ, Schlaggar BL, Petersen SE. Spurious but systematic correlations in functional connectivity MRI networks arise from subject motion. Neuroimage. 2012;59(3):2142–2154. doi:10.1016/j.neuroimage.2011.10.018

40. Nichols TE, Holmes AP. Nonparametric Permutation Tests For Functional Neuroimaging: A Primer with Examples. Hum Brain Mapp. 2001;15:1–25.

41. Rolls ET, Joliot M, Tzourio-Mazoyer N. Implementation of a new parcellation of the orbitofrontal cortex in the automated anatomical labeling atlas. Neuroimage. 2015;122:1–5. doi:10.1016/j.neuroimage.2015.07.075

42. Behrens TEJ, Johansen-Berg H, Woolrich MW, et al. Non-invasive mapping of connections between human thalamus and cortex using diffusion imaging. Nat Neurosci. 2003;6(7):750–757. doi:10.1038/nn1075

43. Behrens TEJ, Woolrich MW, Jenkinson M, et al. Characterization and Propagation of Uncertainty in Diffusion-Weighted MR Imaging. Magn Reson Med. 2003;50(5):1077–1088. doi:10.1002/mrm.10609

44. Kumar VJ, van Oort E, Scheffler K, Beckmann CF, Grodd W. Functional anatomy of the human thalamus at rest. Neuroimage. 2017;147(October 2016):678–691. doi:10.1016/j.neuroimage.2016.12.071

45. Desikan RS, Ségonne F, Fischl B, et al. An automated labeling system for subdividing the human cerebral cortex on MRI scans into gyral based regions of interest. Neuroimage. 2006;31(3):968–980. doi:10.1016/j.neuroimage.2006.01.021

46. Yeh F-C, Panesar S, Fernandes D, et al. Population-Averaged Atlas of the Macroscale Human Structural Connectome and Its Network Topology. Neuroimage. 2018;178:57–68. doi:doi:10.1016/j.neuroimage.2018.05.027.

47. Sperber C, Karnath HO. Impact of correction factors in human brain lesion-behavior inference. Hum Brain Mapp. 2017;38(3):1692–1701. doi:10.1002/hbm.23490

48. Pérennou DA, Leblond C, Amblard B, Micallef JP, Rouget E, Pélissier J. The polymodal sensory cortex is crucial for controlling lateral postural stability: Evidence from stroke patients. Brain Res Bull. 2000;53(3):359–365. doi:10.1016/S0361-9230(00)00360-9

49. Fiori F, Candidi M, Acciarino A, David N, Aglioti SM. The right temporoparietal junction plays a causal role in maintaining the internal representation of verticality. J Neurophysiol. 2015;114(5):2983–2990. doi:10.1152/jn.00289.2015

50. Dieterich M, Brandt T. Perception of verticality and vestibular disorders of balance and falls. Front Neurol. 2019;10(APR):1–15. doi:10.3389/fneur.2019.00172

51. Jahn K, Wagner J, Deutschländer A, et al. Human hippocampal activation during stance and locomotion: FMRI study on healthy, blind, and vestibular-loss subjects. Ann N Y Acad Sci. 2009;1164:229–235. doi:10.1111/j.1749-6632.2009.03770.x

52. Kahane P, Hoffmann D, Minotti L, Berthoz A. Reappraisal of the Human Vestibular Cortex by Cortical Electrical Stimulation Study. Ann Neurol. 2003;54(5):615–624. doi:10.1002/ana.10726

53. Mittelstaedt H. Origin and Processing of Postural Information. 1998;22(4):473–478.

54. Narayanan JT, Watts R, Haddad N, Labar DR, Li PM, Filippi CG. Cerebral activation during vagus nerve stimulation: A functional MR study. Epilepsia. 2002;43(12):1509–1514. doi:10.1046/j.1528-1157.2002.16102.x

55. Sperber C, Griffis J, Kasties V. Indirect structural disconnection - symptom mapping. Brain Struct Funct. 2022;(0123456789). doi:10.1007/s00429-022-02559-x

56. Sherman SM. Thalamus plays a central role in ongoing cortical functioning. Nat Neurosci. 2016;19(4):533–541. doi:10.1038/nn.4269

57. Umarova RM, Reisert M, Beier TU, et al. Attention-network specific alterations of structural connectivity in the undamaged white matter in acute neglect. Hum Brain Mapp. 2014;35(9):4678–4692. doi:10.1002/hbm.22503

58. Lunven M, De Schotten MT, Bourlon C, et al. White matter lesional predictors of chronic visual neglect: A longitudinal study. Brain. 2015;138(3):746–760. doi:10.1093/brain/awu389

59. Herbet G, Duffau H. Contribution of the medial eye field network to the voluntary deployment of visuospatial attention. Nat Commun. 2022;13(1). doi:10.1038/s41467-022-28030-3

60. Rosenzopf H, Wiesen D, Basilakos A, et al. Mapping the human praxis network: an investigation of white matter disconnection in limb apraxia of gesture production. Brain Commun. 2022;4(1):1–14. doi:10.1093/braincomms/fcac004

61. Siegel JS, Ramsey LE, Snyder AZ, et al. Disruptions of network connectivity predict impairment in multiple behavioral domains after stroke. Proc Natl Acad Sci US A. 2016;113(30):E4367–E4376. doi:10.1073/pnas.1521083113

62. Klingbeil J, Wawrzyniak M, Stockert A, Karnath HO, Saur D. Hippocampal diaschisis contributes to anosognosia for hemiplegia: Evidence from lesion network-symptom-mapping. Neuroimage. 2020;208(October 2019):116485. doi:10.1016/j.neuroimage.2019.116485

63. Sperber C, Dadashi A. The influence of sample size and arbitrary statistical thresholds in lesion-network mapping. Brain. 2020;143(5):E40. doi:10.1093/brain/awaa094

64. Pini L, Salvalaggio A, De Filippo De Grazia M, Zorzi M, Thiebaut de Schotten M, Corbetta M. A novel stroke lesion network mapping approach: improved accuracy yet still low deficit prediction. Brain Commun. 2021;3(4). doi:10.1093/braincomms/fcab259

65. Boes AD. Lesion network mapping: Where do we go from here? Brain. 2021;144(1). doi:10.1093/brain/awaa350

66. Cohen AL, Soussand L, Corrow SL, Martinaud O, Barton JJS, Fox MD. Looking beyond the face area: Lesion network mapping of prosopagnosia. Brain. 2019;142(12):3975–3990. doi:10.1093/brain/awz332

67. Corp DT, Joutsa J, Darby RR, et al. Network localization of cervical dystonia based on causal brain lesions. Brain. 2019;142(6):1660–1674. doi:10.1093/brain/awz112

