## Supplementary for "Thalamocortical networks involved in Pusher Syndrome"

### Supplementary materials

**Voxel-based lesion-symptom-mapping (VLSM)**

For VLSM, differences between the two groups (*Pusher+/Pusher-*) were calculated using in-house software in the context of the general linear model within each voxel damaged in at least 5 patients. Lesion size was included as covariate of no interest. To control the family wise error rate, null-distribution of maximum t-scores were also obtained by 5,000 random permutations (Rorden et al., 2007). Results were thresholded at p(FWE) < 0.05 at voxel-level and anatomically labeled using the the AICHA atlas (Joliot et al., 2015). To avoid the right hemispheric lesions from weakening left hemispheric lesion-symptom-associations and vice versa, results were obtained separated by hemisphere. VLSM employing non-parametric permutation testing revealed a significant association of pusher syndrome with lesions in the crus posterior of the right capsula interna (Fig. S1). Hemispheric separation led to no locations significantly associated with pusher syndrome in patients with leisons in the left hemisphere.

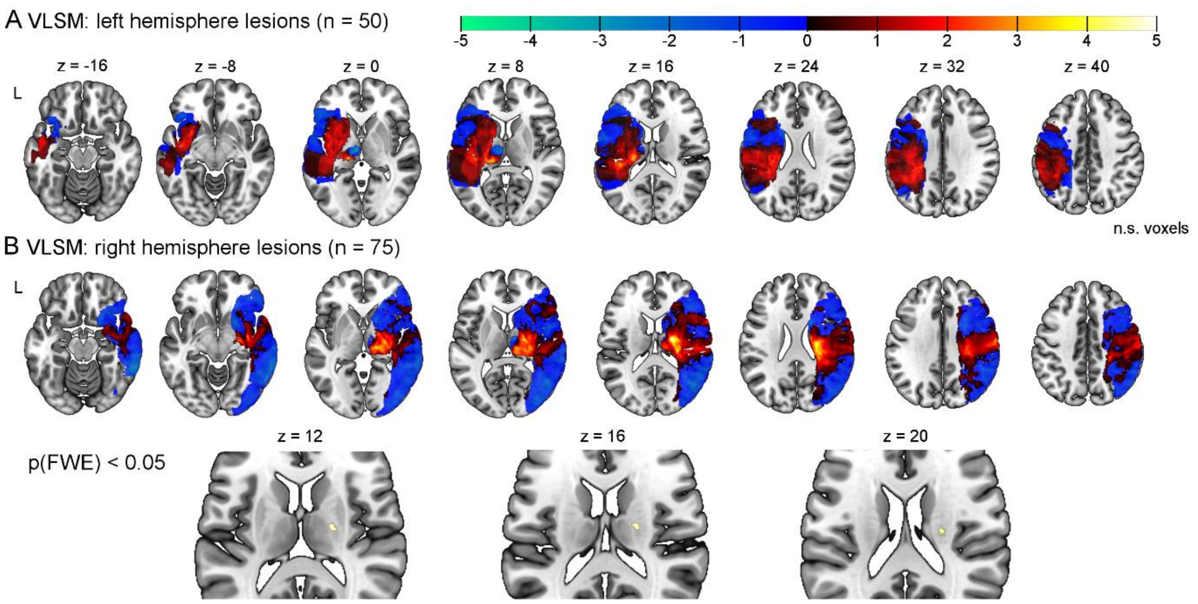


**Figure S1. Voxel-based lesion-symptom-mapping for lesion association with pusher syndrome.** Statistical inference was based on a random permutation test thresholded at p(FWE) < 0.05 at the voxel-level. The VLSM found no significant results for all leftsided lesion (n = 50) (A) but 218 significant voxels in the right capsula interna when restricted to right hemisphere (B). Abbreviations: FWE = family-wise error; L = left. Scale: t-values; coordinates in MNI space.

#### References

Joliot, M., Jobard, G., Naveau, M., Delcroix, N., Petit, L., Zago, L., Crivello, F., Mellet, E., Mazoyer, B., Tzourio-Mazoyer, N., 2015. AICHA: An atlas of intrinsic connectivity of homotopic areas. Journal of neuroscience methods 254, 46–59. https://doi.org/10.1016/j.jneumeth.2015.07.013.

Rorden, C., Karnath, H.-O., Bonilha, L., 2007. Improving lesion-symptom mapping. Journal of cognitive neuroscience 19, 1081–1088. https://doi.org/10.1162/jocn.2007.19.7.1081.
